# Global cooling & the rise of modern grasslands: Revealing cause & effect of environmental change on insect diversification dynamics

**DOI:** 10.1101/392712

**Authors:** Katie E. Davis, Adam T. Bakewell, Jon Hill, Hojun Song, Peter Mayhew

## Abstract

Utilising geo-historical environmental data to disentangle cause and effect in complex natural systems is a major goal in our quest to better understand how climate change has shaped life on Earth. Global temperature is known to drive biotic change over macro-evolutionary time-scales but the mechanisms by which it acts are often unclear. Here, we model speciation rates for Orthoptera within a phylogenetic framework and use this to demonstrate that global cooling is strongly correlated with increased speciation rates. Transfer Entropy analyses reveal the presence of one or more additional processes that are required to explain the information transfer from global temperature to Orthoptera speciation. We identify the rise of C_4_ grasslands as one such mechanism operating from the Miocene onwards. We therefore demonstrate the value of the geological record in increasing our understanding of climate change on macro-evolutionary and macro-ecological processes.

## Introduction

Global environmental change has played a major role through geological time in shaping the diversity of life on Earth we see today (Vermeij, 1978; Vrba, 1980; Benton, 2009; Kozak & Wiens, 2010), and in the face of indisputable climate change, there is increasing focus on how this past change can inform our understanding of the ongoing biodiversity crisis. Increasingly, research is focussing on how we can bridge the gap between palaeontology and ecology to best utilise the information held within the geological record (Lyons & Wagner 2009; Willis et al. 2010; Escarguel et al. 2011; Fritz et al. 2013; Dietl et al. 2014). Palaeobiology and palaeoecology also have much to offer the field of biodiversity informatics, which seeks to answer many of the same questions in understanding the history of life of Earth and how it has been shaped by environmental forces (Peterson et al. 2010).

The geological record bears witness to many instances of climate change and subsequent biotic change and yet remains a largely untapped resource. The Cenozoic Era (66Ma – present) alone has experienced a dramatic shift from greenhouse to icehouse conditions. Recognisably modern fauna and flora arose during the Oligocene and Miocene (Kraatz & Geisler 2010); a period of environmental transition in which previously dominating forest habitats began to fragment with the expansion of modern grassland ecosystems (Strömberg 2005; Edwards et al. 2010; Strömberg 2011). This period was characterised by significant global cooling after the tropical, greenhouse conditions that dominated following the Palaeocene-Eocene Thermal Maximum (PETM) (Mudelsee et al. 2014). This cooling trend continued with a descent into icehouse conditions after the Eocene Optimum, culminating in the onset of Antarctic glaciation at the Eocene-Oligocene transition (Ivany et al. 2000; Barker et al. 2007; Liu et al. 2009). This transitional period also resulted in extinctions and large-scale faunal turnover in marine invertebrates and mammals at the Eocene-Oligocene boundary (Sun et al. 2014). During the Oligocene forests still dominated but C4 grasses began to radiate and open grassland habitats became more widespread (Strömberg 2005; Edwards et al. 2010; Strömberg 2011). Fully modern ecosystems originated following the Middle Miocene Disruption (~14 Ma); a period characterised by further global cooling, and the widespread replacement of forests with open grassland (Strömberg 2005). Fully modern C_4_ grasslands became established between 3–8 Ma (Edwards et al. 2010). These environmental shifts were accompanied by major changes to the terrestrial fauna; most notably the mammalian fauna, which radiated to take advantage of the widespread grassland habitats (Cerling et al. 1998; Codron et al. 1998; Janis et al. 2002; Bobe & Behrensmeyer 2004). Recent research suggests that climate change and the opening up of grassland habitats at this time may also have played a role in shaping insect evolution (e.g., Peña & Wahlberg 2008; Voje et al. 2009; Toussaint et al. 2012; Lo et al. 2017).

The development of modern numerical modelling techniques have now been utilised to demonstrate a correlation between environmental change and diversification within a statistical framework (e.g., Figueirido et al. 2011; Martin et al. 2014; Claramunt & Cracraft 2015; Mannion et al. 2015; Davis et al. 2016). Showing a correlation, however, does not necessarily show causation and the mechanisms by which climate change affects biological diversification can be more difficult to elucidate, though various explanations have been proposed including habitat fragmentation, new habitat availability and adaptive radiation (e.g., Peña & Wahlberg 2008; Voje et al. 2009; Claramunt & Cracraft 2015; Davis et al. 2016; Davis et al. 2018). Information Theory is a statistical tool that has been successfully utilised to identify cause and effect between time series in geological and palaeontological data sets (e.g., Dunhill et al. 2014; Liow et al. 2015). The idea of using time series as a tool to predict relationships was first proposed in 1956 (Wiener 1956), whilst Hannisdal (2011) was the first to introduce the use of information theory to detect directionality between palaeontological time series. These methods have been primarily developed for short, irregularly spaced data series; where continuous time series are available, such as those obtained from environmental proxy time series and speciation rate curves, it is possible to utilise a measure that makes fewer assumptions of the data series. Transfer Entropy (TE) (Schreiber 2000) is a type of Information Theory that does not assume linearity in the linking process; it is therefore a powerful method in cases where the exact linking processes are unknown.

The insect order Orthoptera includes grasshoppers, locusts, crickets, katydids and wetas; they inhabit a diverse array of habitats from tropical rainforest to grasslands to desert. The first definitive orthopteran fossil is the 300 million year old *Oedischia williamsoni* from the Permian of France (Song et al. 2015). Further fossil evidence suggests that the two suborders Ensifera (crickets, katydids, weta) and Caelifera (grasshoppers) had diverged by around 260 Ma but little is known about how Orthoptera diversified to produce 27,000 extant species or the processes driving their evolution. As a species-rich clade with a long geological history there are undoubtedly a variety of factors that have shaped their extant diversity but here, we ask whether past environmental change had an impact on orthopteran diversification. In particular, we ask whether the Miocene origin of modern grasslands drove adaptive radiation in grassland species. We achieve this by using supertree methods to build a new time-calibrated phylogenetic hypothesis for Orthoptera. We use this phylogeny to explore orthopteran diversification dynamics through time then test for causality between past environmental change and speciation rates using Transfer Entropy. We find that speciation rates in Orthoptera are strongly correlated with global cooling and that significant increases in net diversification occurred during global cooling events and coincident with the origin and spread of C4 grasslands. Trait-based analyses show that speciation rates recovered for species associated with grasslands and other open-habitats are an order of magnitude higher than those of species found in forests and other habitats. Considering just the last 17 Ma (Miocene – Recent) we find that speciation rates are strongly correlated with both global cooling and C4 grassland expansion. Our Transfer Entropy analyses reveal that over this period, grassland expansion is the mechanism via which global cooling stimulated biological diversification.

## Methods

### Data collection and processing

The Web of Knowledge Science Citation Index (wok.mimas.org) was used to identify all papers containing, or potentially containing, phylogenetic trees for Orthoptera. The years 1980–2015 were searched using the search terms: phylog*, taxonom*, systematic*, divers*, cryptic and clad* in conjunction with all scientific and common names for Orthoptera from sub-order to family level. All source trees and meta-data were digitised in their published form using TreeView (Page 1996) and the Supertree Toolkit (STK – Davis & Hill 2010; Hill & Davis 2014). Along with the tree string in Newick format, meta-data were stored including: bibliographic information, character type, and phylogenetic inference. No corrections were made for synonyms or any other apparent errors or inconsistencies in the source trees at this stage. See Supporting Appendices SA1 and SA2 for source trees and source tree references.

Data processing prior to supertree construction was carried out in a consistent and clearly documented manner. We followed the protocol as previously used in other supertree analyses (e.g., Davis & Page 2014; Davis et al. 2015) and outlined here as follows. All included phylogenies were required to fulfil three criteria before inclusion in the data set: 1) be presented explicitly as a reconstruction of evolutionary relationships; 2) be comprised of clearly identifiable species, genera or higher taxa and clearly identifiable characters; 3) be derived from the analysis of a novel, independent dataset. All taxon names, including higher taxa, were standardised following the Orthoptera Species File (Cigliano et al. 2018). Taxonomic overlap was set such that each source tree was required to have a minimum of at least two taxa in common with at least one other source tree (Sanderson et al. 1998). The data set did not satisfy the overlap requirements and after removal of “islands” of unconnected source trees the taxa number was reduced from 2,258 to 1,748.

### Supertree construction

Despite ongoing active development of new methods (Akanni et al. 2014; Oliveira Martins et al. 2016), Matrix Representation with Parsimony (MRP – Baum & Ragan 2004) is still the most tractable supertree method for large datasets. We therefore utilised MRP to infer a phylogenetic supertree from a total data set of 261 source trees taken from 124 papers published between 1991 and 2015. A down-weighted (weight=0.1) taxonomy tree was included to improve performance and data cohesion (Bininda-Emonds & Sanderson 2001). Source trees were encoded as a series of group inclusion characters using standard Baum and Ragan coding (Baum & Ragan 2004), and automated within the STK software (Davis & Hill 2010; Hill & Davis 2014). All taxa subtended by a given node in a source tree were scored as “1”, taxa not subtended from that node were scored as “0”, and taxa not present in that source tree were scored as “?”. Trees were rooted with a hypothetical, “all zero” outgroup. Large taxon numbers greatly increase computational time and reduce chances of finding the shortest trees, therefore the data set was split into two partitions representing the two reciprocally monophyletic suborders: Caelifera (1,377 included species) and Ensifera (789 included species). The resulting MRP matrices were analysed using standard parsimony algorithms in TNT (Goloboff et al. 2008). We used the “xmult=10” option, and ran 1,000 replicates for the analysis, each using a different random starting point for the heuristic search. This improved exploratory coverage of the tree space, potentially avoiding local minima in the solutions. We computed a Maximum Agreement Subtree (MAST) using PAUP* (Swofford 2002) to remove conflicting leaves, reducing the total number across both data partitions from 1,748 to 1,293. Although not limited to supertree methods, one disadvantage of the MRP method is that it can lead to the creation of spurious clades and relationships that are not present in any of the source trees (Bininda-Emonds & Bryant 1998; Davis & Page 2014). These misplaced taxa are generally referred to as “rogue taxa” and are usually a result of either poorly constrained or poorly represented taxa within the source trees. Studies have shown that identifying and removing rogue taxa *a priori* can create further problems, as rogues still have the potential to phylogenetically constrain the positions of other taxa (Trautwein et al. 2011). Hence *a priori* removal often creates new rogue taxa. We identified a small number of rogue taxa (~2%) in the resulting tree. It is important that these novel clades are not interpreted as biologically meaningful and therefore should be removed before undertaking further analysis (Pisani & Wilkinson, 2002). We provide a list of removed taxa in Appendix SA3. To maximise taxon coverage, an additional 226 taxa were added to the tree based on their positions in source trees excluded at the data overlap stage (see Appendices SA4 and SA5 for source trees and references). This resulted in a final supertree comprising 1,519 taxa, as compared to the previous largest orthopteran phylogeny that contained 274 taxa (Song et al. 2015).

### Supertree time-calibration

Phylogenies derived from parsimony analyses do not have meaningful branch lengths that can be used to infer the absolute diversification times and rates in the tree, they can only inform on the relative times of divergence. It is necessary, therefore, to use external reference points to time-calibrate trees inferred using parsimony. We used a combination of fossil calibrations supplemented by biogeographic calibrations to time-scale our tree. Forty-three nodes were calibrated using fossil first occurrence data downloaded from Fossilworks (Data available from the Fossilworks database: fossilworks.org). These fossils were assigned phylogenetically to either the stem or crown of clades using the taxonomy assigned in Fossilworks. An additional seven geological calibration points were obtained from published molecular phylogenetic analyses (see Appendix ST1 for node numbers and calibration dates and Appendix SF1 for the tree with the calibrated nodes labelled as in the CSV file). The R package “paleotree” (Bapst 2012) was used to scale the tree and to extrapolate dates to the remaining nodes. To extend node calibrations to the whole tree, we used the “equal” method, with minimum branch lengths set to 0.1 Myr. See Appendix SA6 for the time-calibrated supertree.

### Environmental correlations

Correlation analyses were carried out via DCCA: detrended cross-correlation analysis, which is designed for correlating non-stationary time series (Kristoufek 2014). Speciation rate time series were correlated against both palaeo-temperature and palaeo-vegetation to assess whether a statistically significant correlation exists. All analyses were carried out in R 3.2.2 (R Core Team 2017).

Speciation rate curves were obtained from BAMM (Bayesian Analysis of Macroevolutionary Mixtures), which implements a Metropolis Coupled Monte Carlo (MCMC) approach to calculate diversification rates and significant rate shifts along lineages (Rabosky 2014). Four chains were executed for the analysis, each with a total of 30 million generations executed, with a minimum clade size of five taxa used to aid convergence. Ten thousand of the results were stored, with 1,000 discarded as “burn-in”, leaving 9,000 samples for subsequent analysis with regards to temperature correlation. The analysis also accounted for non-complete coverage of taxa in the tree by specifying a clade-dependent sampling bias factor derived from the taxonomy in Orthoptera Species File (Cigliano et al. 2018).

The palaeo-temperature data were obtained from oxygen isotope (δ^18^O) records (Veizer et al. 1999; Zachos 2001) and the δ^18^O curve was first smoothed to remove autocorrelation. The speciation rate curves from BAMM consist of 9,000 individual curves (10,000 minus 10% burn in), therefore a total of 9,000 correlations per test were carried out with each DCCA coefficient saved. The saved DCCA coefficients were then plotted as a distribution and a Wilcoxon rank sum test was used to test if the mean of the distribution was non-zero. Global palaeo-temperature correlations were carried out for both the full evolutionary history of Orthoptera and also partitioned at 17 Ma to allow direct comparison with the palaeo-vegetation time series.

Palaeo-vegetation data for the last 17 Ma were extracted from Osborne (2008). Three regions were considered: Pakistan, the Great Plains (USA) and East Africa. The first two used palaeosol data, whilst the latter used tooth enamel to obtain δ^13^C; where higher δ^13^C is a proxy for increased C_4_ biomass. After checking location data held in GBIF for the species in our tree we discarded the East Africa and Pakistan data as this showed a significant geographic bias towards Europe and the USA with very few species from East Africa and Pakistan represented (Appendix SF2). In order to obtain a continuous time series a cubic smooth spline was used to interpolate the point data. These time series were then correlated against the 9,000 BAMM simulations, and plotted as a distribution, in the same manner as for the palaeo-temperature correlations.

### Temporal-diversification analyses

TreePar (Stadler 2011) was used to assess changes in net diversification rates across the tree through time. The “bd.shifts.optim” function was used, together with a 1 million year grid, allowing rate changes to be assessed at 1 million year intervals. The analyses were run across the whole tree starting from the present day, back to the root node. Net diversification rates were allowed to be negative and we set TreePar to look for up to 10 temporal changes in net diversification rate.

### Trait-diversification analyses

MUSSE (MultiState Speciation and Extinction) was implemented in Diversitree (Fitzjohn 2012) to model diversification rates based on habitat trait data. Broadly defined habitat types for as many species as possible were collected via an exhaustive literature search (Appendix ST2). Taxa were designated as having habitat preferences as follows: “open”, “closed”, or “mixed”; where open = grassland, desert etc, closed = forest, woodlands etc, and mixed = found in both open and closed habitats. Habitat trait data were obtained for 377 out of 1,519 species in the phylogeny, the remainder were coded as “NA”.

### Information transfer

Transfer entropy (TE) is a directional information flow method that quantifies the coherence between continuous variables in time (Schreiber 2000). It is an extension of the mutual information method, but can take into account the direction of information transfer using an assumption that the processes can be described by a Markov model. Transfer Entropy reduces to a linear Granger causality process, whereby a signal in one time series gives a linear response to the second time series, when the two time series can be linked via autoregressive processes (Granger 1969; Amblard & Michel 2012). However, TE makes fewer assumptions on the linearity of the processes involved and hence is more suitable for analysing causality when the processes involved are unknown (Lungarella et al. 2007; Ver Steeg & Galstyan 2012). Transfer Entropy is calculated using:

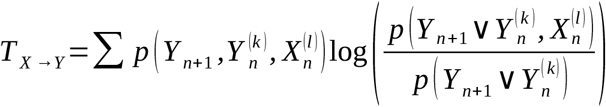

where T_x→y_ is the TE from time series *X* to time series *Y*, both of which have data at time *n*, and *k* and *l* are the embedding dimensions of the two time series respectively. We used the R (R Core Team 2017) package “TransferEntropy” (Torbati & Lawyer 2016) which implements the above equation using a nearest neighbour algorithm (Kraskov et al. 2003). This function returns a numeric value where 0 indicates no information transfer, positive numbers indicate information transfer, and negative numbers indicate misinformation transfer which indicates the presence of other processes in operation (Bossomaier et al. 2016). The embedding dimensions of the time series were estimated using the R package “nonlinearTseries” (Garcia & Sawitzki 2015).

We calculated TE for two time series over the full tree: speciation rate and δ^18^O, and for three time series for the last 17 Ma: speciation rate, δ^18^O and δ^13^C; where δ^18^O represents the palaeo-temperature proxy and δ^13^C represents the palaeo-vegetation proxy. We therefore derived two TE values for the full tree - speciation – δ^18^O, and vice versa; and six TE values for the truncated Miocene data: speciation - δ^18^O, speciation - δ^13^C and δ^18^O - δ^13^C, in both directions of transfer. Significance was tested by creating 250 surrogate time series by randomising the “source” time series, leaving the “target” time series unchanged. If the transfer entropy value was outside the 95% interval of the surrogate transfer entropy values it was deemed significant (Chávez et al. 2003).

## Results

### Supertree construction

Our final supertree (Fig. 1) contained 1,519 taxa and is broadly consistent with recent orthopteran phylogenies (e.g., Flook et al. 1999; Sheffield et al. 2010; Zhou et al. 2010; Song et al. 2015). All 15 superfamilies are represented and 41 out of 44 families, as defined by the Orthoptera Species File (Cigliano et al. 2018). All superfamilies were recovered as monophyletic. Families were largely recovered as monophyletic with the notable exception of Romaleidae and Dericorythidae, which are split and nested within Acrididae. This finding is, however, reflected in the source data (e.g., Li et al. 2011; Song et al. 2015).

**Figure 1:**
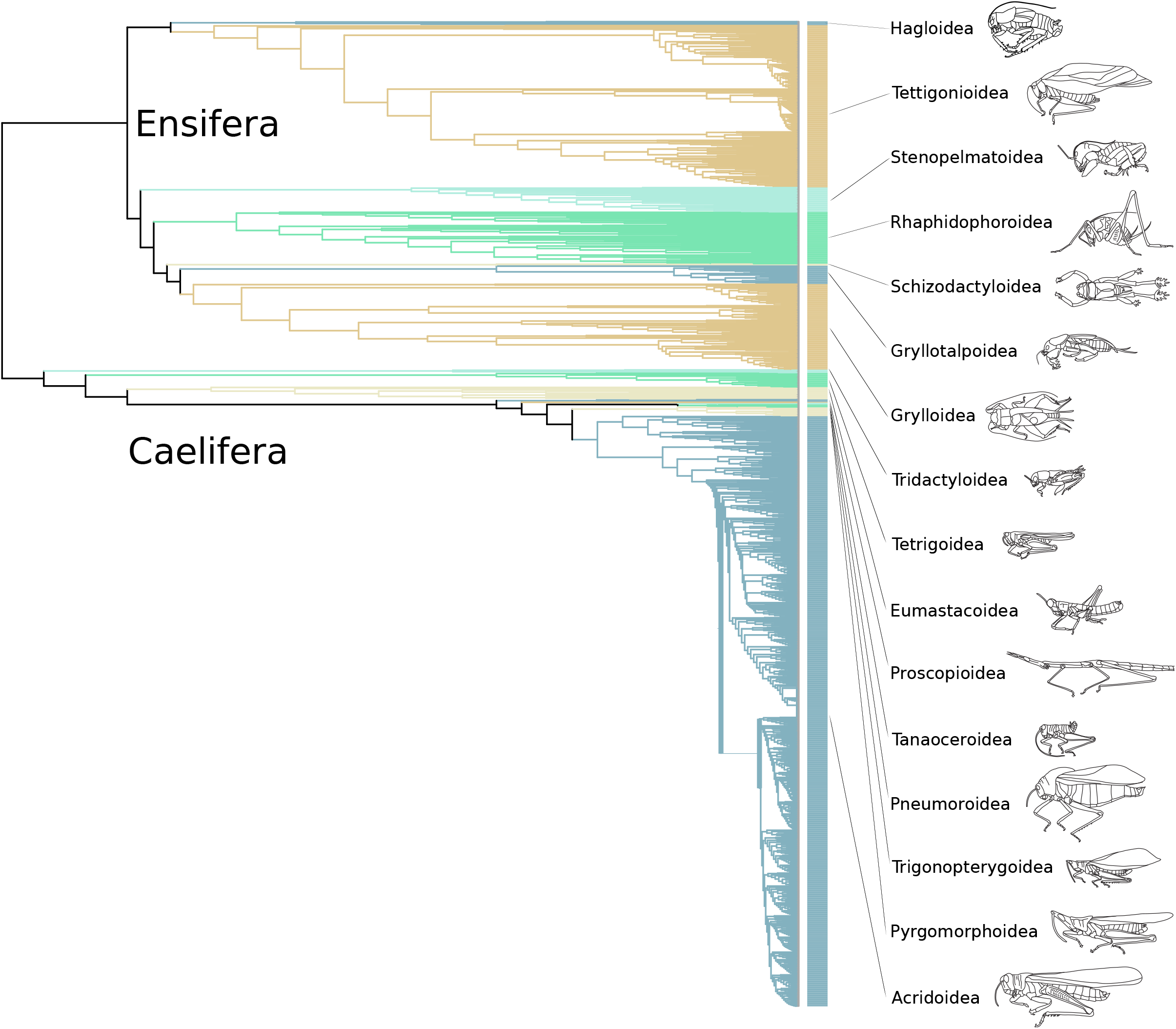
Phylogenetic supertree of Orthoptera Maximum Agreement Subtree (MAST) from MRP analyses shown. Superfamilies are labelled on the right hand side along with representative line drawings (line drawings from Song *et al.* 2015).

### Environmental correlations

Using the 9,000 sets of speciation rate time series extracted from BAMM for the full tree (Rabosky 2014; Rabosky et al. 2014) we found that speciation rate was strongly correlated with global palaeo-temperature (Fig. 2). For each set of extracted speciation rates a correlation coefficient was calculated between -1 (speciation rate increases with cooler temperatures) and 1 (speciation rate increases with warmer temperature). For a correlation coefficient of zero, temperature has no effect on speciation rate. The distribution was not normally distributed, therefore we used a Wilcoxon signed rank test to test whether the distribution of all 9,000 correlation coefficients differed from the null hypothesis of a zero mean correlation coefficient (i.e. no temperature correlation). We found a strong negative mean correlation of r = -0.3801 (SD = 0.0161, p < 2.2e-16 detrended cross-correlation analysis). For the truncated 17 Ma time series we also found a strong negative correlation between speciation rate and global temperature of r = -0.4220 (SD =0.0094, p < 2.2e-16 detrended cross-correlation analysis) (Fig. 2).

**Figure 2:**
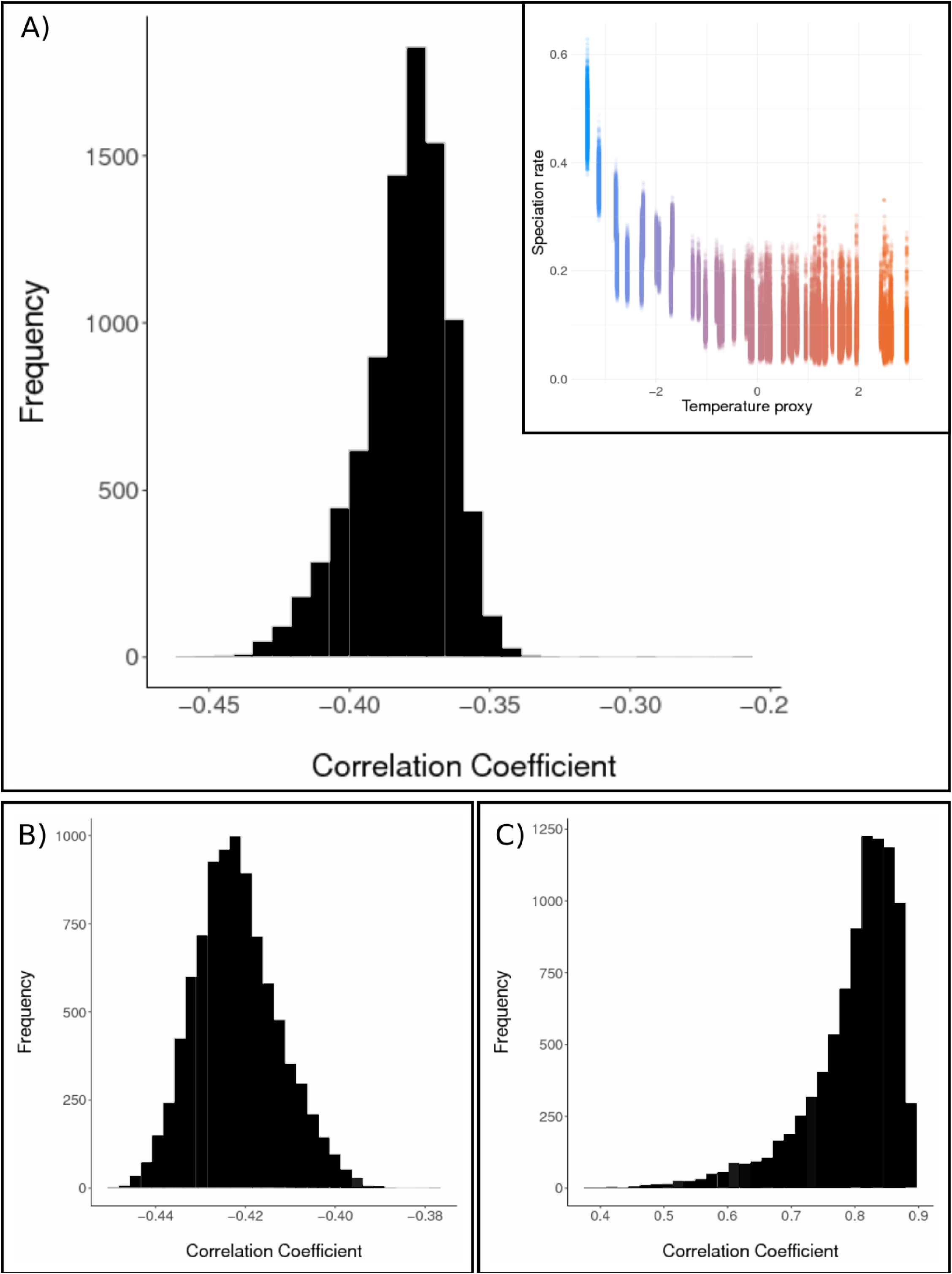
Speciation rates, extracted from BAMM (Rabsoky *et al.* 2014) plotted against environmental correlations. Top panel (A): main figure – histogram showing the frequency distribution of correlation coefficients between speciation rate and global temperature proxy for the whole tree, showing that there is a negative correlation between speciation rates and temperature; inset – speciation rates plotted against global temperature proxy, showing that speciation rates increases as global temperatures decrease, blue represents cooler temperatures and red represents warmer temperatures. Bottom panel: (B) histogram showing the frequency distribution of correlation coefficients between speciation rate and global temperature proxy for the 17 Ma – Recent partition, showing that there is a negative correlation between speciation rates and temperature; (C) histogram showing the frequency distribution of correlation coefficients between speciation rate and the C_4_ biomass proxy for the 17 Ma – Recent partition, showing that there is a positive correlation between speciation rates and C_4_ biomass.

The palaeo-vegetation correlation analyses using the US vegetation data (Osborne 2008) also utilised the truncated speciation rate time series. For these, the correlation coefficient was calculated between -1 (speciation rate decreases with C4 biomass) and 1 (speciation rate increases with C4 biomass). For a correlation coefficient of zero, grassland abundance has no effect on speciation rate. Again, the distributions recovered were not normally distributed; we therefore used a Wilcoxon signed rank test to test whether the distribution of all 9,000 correlation coefficients differed from the null hypothesis of a zero mean correlation coefficient (i.e. no grassland abundance correlation). We found a very strong positive correlation of r = 0.7972 (SD = 0.0736, p < 2.2e-16 detrended cross-correlation analysis) (Fig. 2).

### Temporal-diversification analyses

Our TreePar (Stadler 2011) analyses rejected a constant rate diversification model. The best fit model, as identified by AICc and LRT (AICc=11602.76, p=0.0003), recovered seven rate shifts (Table 1). A model allowing eight shifts did not significantly improve the likelihood. Three of these shifts represent increases in net diversification rate and correspond directly to the following global cooling events (Fig 3); the Albian-Aptian “cold snap” (113 Ma – Mutterlose et al. 2009), the Eocene-Oligocene transition (34 Ma – Liu et al. 2009), and the Middle Miocene disruption (12 Ma – Shevenell et al. 2004). The last of these also corresponds to a burst of C4 grassland evolution (Osborne 2008; Edwards et al. 2010), whilst the Eocene-Oligocene transition is closely coincident with the origins of C4 grasses (Edwards et al. 2010). Finally, we detect a decrease in net diversification rate at the Cretaceous-Palaeogene boundary (66 Ma).

**Table 1.** Results from TreePar analysis. For each model the AIC and corrected AIC were calculated. The lowest AICc was for eight rate shifts, but this was not significantly different from seven shifts highlighted), which we therefore consider the optimum model. All significant AICc are **bold**.

| Model | 0 shifts | 1 shift | 2 shifts | 3 shifts | 4 shifts | 5 shifts | 6 shifts | 7 shifts | 8 shifts | 9 shifts | 10 shifts |
| --- | --- | --- | --- | --- | --- | --- | --- | --- | --- | --- | --- |
| logL | 5855.483 | 5820.809 | 5808.46 | 5802.315 | 5796.107 | 5792.356 | 5785.832 | 5778.008 | 5773.815 | 5770.954 | 5768.563 |
| AIC | 11714.97 | 11651.62 | 11632.92 | 11626.63 | 11620.21 | 11618.71 | 11611.66 | 11602.02 | 11599.63 | 11599.91 | 11601.13 |
| AICc | 11714.97 | **11651.66** | **11633.02** | **11626.81** | **11620.49** | 11619.12 | 11612.22 | **11602.76** | 11600.57 | 11601.08 | 11602.55 |

**Figure 3:**
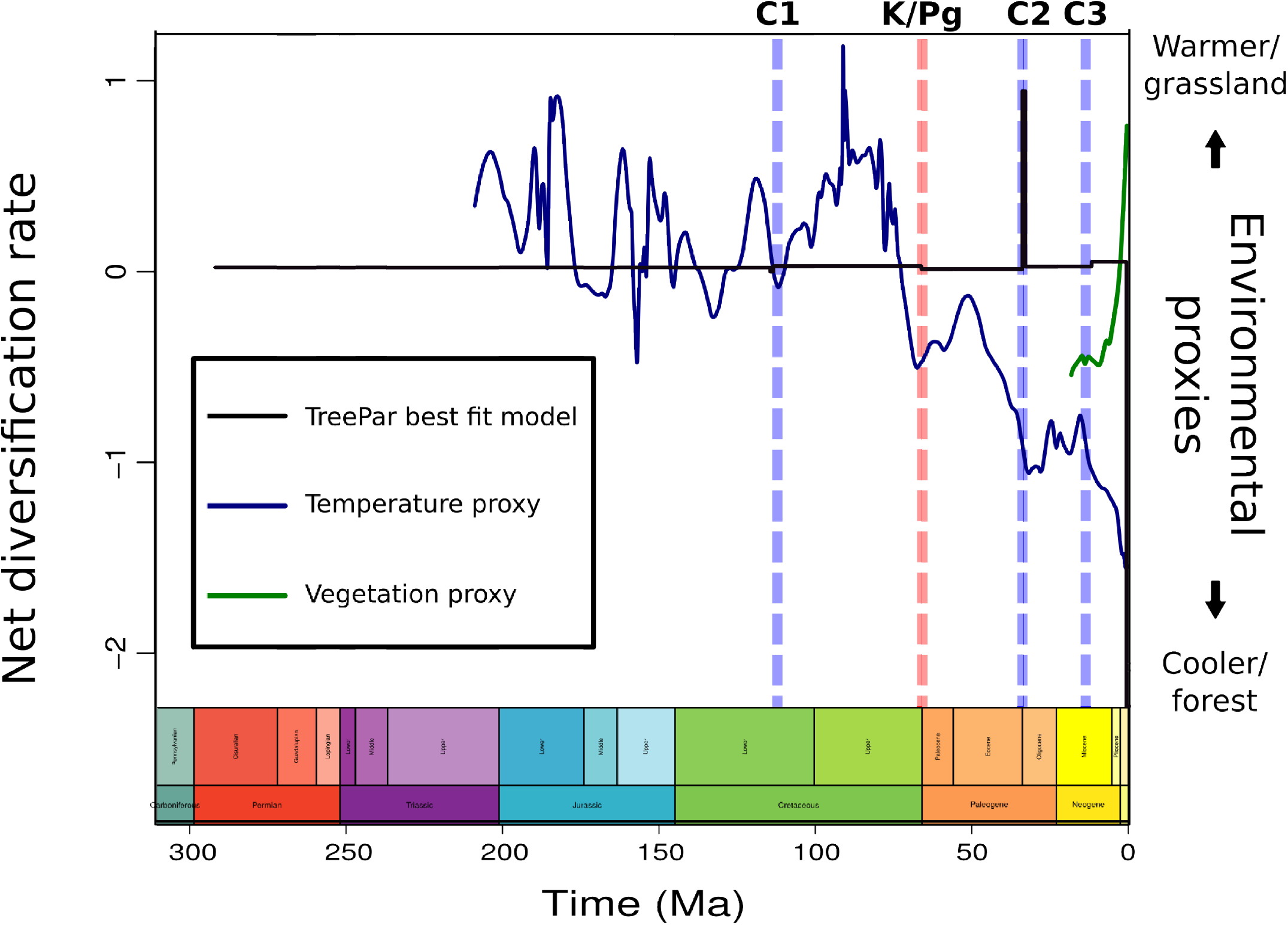
TreePar best fit model of rate shifts (black line) plotted against palaeo-temperature (blue line) and palaeo-vegetation (green line) and scaled to geological time. Geological time scale was added using the R package “strap” (Bell & Lloyd 2015). Global events labelled as C1: Albian-Aptian cold snap; C2 = Eocene-Oligocene transition; C3 = Middle Miocene disruption; K/Pg = Cretaceous-Palaeogene mass extinction event.

### Trait-diversification analyses

Trait-based analyses, implemented in MUSSE (FitzJohn 2012) rejected a null model whereby speciation, extinction and net diversification rates are unrelated to habitat (Fig. 4). Our analyses found that speciation rates in open habitats were an order of magnitude higher than those found for closed or mixed habitats with no overlap of the confidence intervals (open: mean = 3.5065, SD = 1.541988; closed: mean = 0.3906, SD = 0.0246, mixed: mean = 0.4991, SD = 0.2657). Extinction rates reveal a similar pattern though the 2.5% confidence interval for open habitats overlaps with those for closed and mixed habitats (open: mean = 3.028975, SD = 1.618406; closed: mean = 0.3696197, SD = 0.02499524, mixed: mean = 0.308759, SD = 0.2997). Net diversification reveals highest rates in open habitats, with an overlap of the 2.5% confidence interval with mixed habitats. Closed habitats have the lowest net diversification rates (open: mean = 0.4775, SD = 0.0866; closed: mean = 0.0210, SD = 0.0017, mixed: mean = 0.1903, SD = 0.0397).

**Figure 4:**
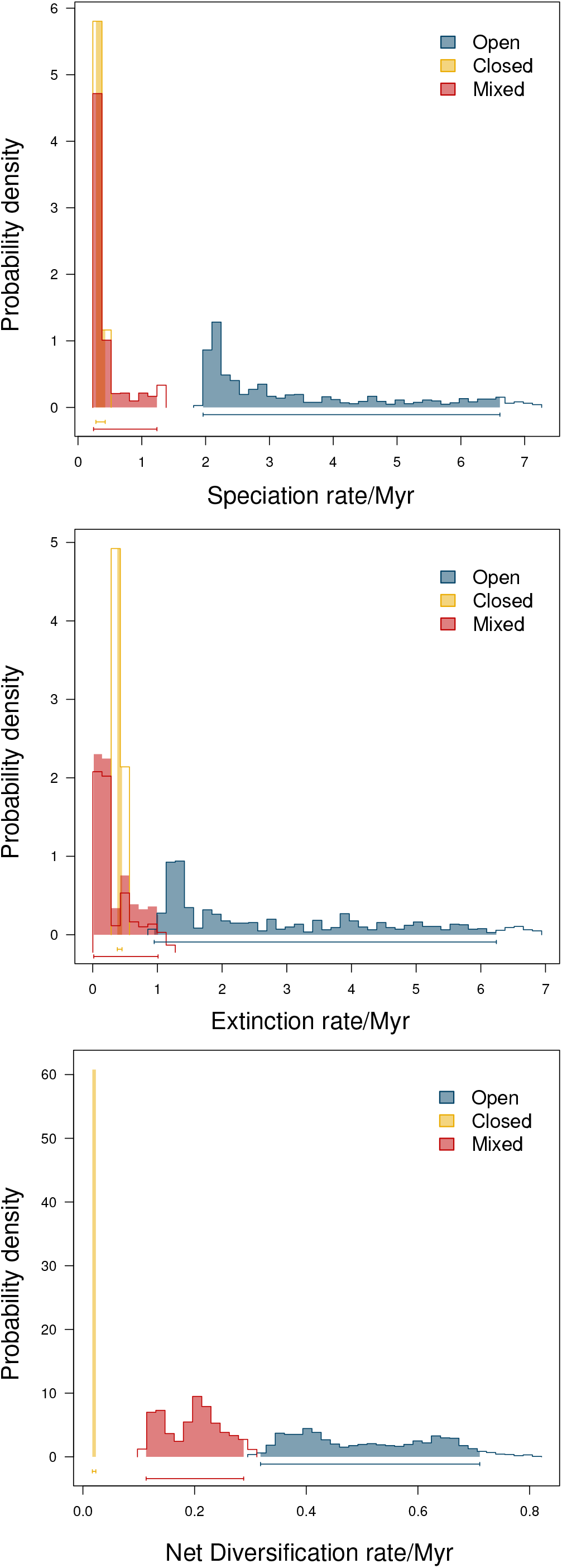
Trait diversification rates. Rates modelled in MUSSE (Fitzjohn 2012) and probability density plotted for speciation rate (top panel), extinction rate (middle panel) and net diversification rate (bottom panel). Blue = open habitat-associated species; yellow = closed habitats; red = mixed habitat associations.

### Information transfer

The results from the TE analysis for the full tree shows a mis-information signal (negative value for TE) from temperature to speciation rate (-0.7485), indicating the presence of one or more hidden drivers (Bossomaier et al. 2016). There is no information transfer from speciation rate to temperature (-0.0028). The Miocene data set again shows a flow of mis-information from temperature to speciation rate (-0.1134) with no information flow in the reverse direction (-0.0522). When the palaeo-vegetation data are introduced we recover information flow from temperature to vegetation (0.1359) with a mis-information signal from vegetation to temperature (-0.1254). We also recover a signal from vegetation to speciation rate (0.1697) but no signal in the reverse direction (-0.004). All the Transfer Entropy results were significant at the 95% confidence interval (Table 2).

**Table 2.** Results of the Transfer Entropy analysis. For each of the three time periods used, the TE score is given along with the 95% confidence interval in parentheses below. Significant results in **bold**. V: Vegetation proxy, S: Speciation rate, T: Temperature proxy. Arrows indication direction of information flow.

| Time period | S → T | T → S | V → T | T → V | S → V | V → S |
| --- | --- | --- | --- | --- | --- | --- |
| 17Ma → 0ma | -0.052<br>(-0.039 - 0.025) | -0.113<br>(-3.009 - -2.917) | -0.125<br>(-0.514 - -0.198) | 0.136<br>(-0.266 - 0.122) | -0.004<br>(-0.002 - 0.227) | 0.170<br>(-1.609 - -1.245) |
| 220Ma → 0Ma | -0.003<br>(-0.021 - 0.016) | -0.749<br>(-3.006 - -2.914) |  |  |  |  |

### Discussion

Disentangling past cause and effect in complex natural systems through geological time is a major goal in palaeobiology. The use of the geological record to inform our understanding of the effects of present day climate change on biodiversity is an ever-growing field of research but if palaeobiologists intend to make an impact and contribution to conservation efforts, it is vitally important that we are able to describe not only the *how* and *when,* ie. the nature of the relationship between large-scale biotic and abiotic changes through Earth’s history but also to address *why* these changes arose.

When it comes to the *how* and *when* it is widely recognised that global climate change has had an impact on the biota over geological time scales (Vermeij 1978; Vrba 1980; Benton 2009; Kozak & Wiens 2010). Mean global temperature, in particular, has been demonstrated to show a strong correlation with diversification rates in taxa as diverse as hermit crabs and squat lobsters (Davis et al. 2016), crocodiles (Mannion et al. 2015), and birds (Claramunt & Cracraft 2015). Erwin (2009) suggested that the positive association between biodiversity and global temperature at the spatial scale would predict a positive relationship between biodiversity and global temperatures temporally. While this has been supported by some studies (e.g., Figueirido et al. 2011; Mannion et al. 2015), others have found the opposite (Claramunt & Cracraft 2015); Davis et al. 2016) or even no relationship (Mannion et al. 2015). Identifying the *why,* ie. the underlying mechanisms by which climate change drives biotic turnover is more difficult but the cross-field application of Information Theory allows us to test for correlations between ecological and palaeontological time series (e.g., Dunhill et al. 2014; Liow et al. 2015).

In this paper we show *how* climate change has impacted on biological diversification in orthopteran insects through geological time. Our analyses reveal a consistent pattern in which global cooling through geological time contributed to shaping the evolutionary history of Orthoptera. Two different models – clade and temporal - of diversification dynamics support this, revealing a picture in which speciation rates are correlated with global cooling and that significant increases in net diversification occurred during global cooling events. For the Miocene to Recent we are also able to demonstrate *why* climate change impacted upon speciation rates in Orthoptera, by showing that vegetation change (in the form of increased C_4_ biomass) was one mechanism by which global cooling drove speciation in Orthoptera. We suggest that this took place via forest fragmentation and the expansion of open C_4_ grasslands providing vacant niche space into which adaptive radiation could take place in the absence of competition. This is congruent with the presence of an increase in net diversification rate at 12 Ma, coincident with the Middle Miocene disruption cooling event and with palaeobotanical evidence for a burst of C_4_ grassland evolution (Osborne 2008; Edwards et al. 2010). Our trait analyses further support this by revealing that species living in open habitats have significantly higher speciation rates than species living in closed or mixed habitats.

At present, the underlying mechanism(s) by which global cooling promotes speciation through deeper geological time remain unknown, though we conjecture that climate change induced habitat change is a plausible explanation. The largest increase in net diversification was found at the Eocene-Oligocene transition (34 Ma), a period defined by the onset of Antarctic glaciation (Pound & Salzmann 2017). Phytolith assemblages from the central Great Plains indicate that open habitat C3 grasses first migrated into subtropical, closed forest at this time (Strömberg 2011). The trigger for this opening up and expansion of grasslands is not well understood (Strömberg 2011) but we suggest that global cooling driven habitat fragmentation may again be one possible mechanism. The further back in time we look, the harder it becomes to explain the observed patterns and the earliest evidence we have for an increase in net diversification took place during the Cretaceous. Though widely held to have experienced a consistent greenhouse climate, evidence from sedimentology reveals a brief icehouse interlude in the mid-Cretaceous – the Albian-Aptian “cold snap” (Mutterlose et al. 2009), at which our first diversification increase is exactly coincident (113 Ma). The Cretaceous was also a time of major biotic turnover and ecosystem reorganisation, known as the Cretaceous Terrestrial Revolution (KTR) in which the previously dominant conifererous forests were rapidly replaced by angiosperms (Lloyd et al. 2008). This period, dating to 125–80 Ma, is characterised by a burst of diversification in angiosperms, insects, reptiles, birds and mammals. It is therefore possible that the signal recovered by our analysis at 113 Ma represents the impact of the KTR on orthopteran diversification. Though we present evidence indicating that global cooling driven forest fragmentation and the expansion of open grasslands may be one plausible driver of speciation throughout their evolutionary history, it is important to note that Orthoptera are an old, diverse group, and geological processes such as plate tectonics and orogenesis, as well as evolutionary processes such as sexual selection acting on acoustic signal are likely to have played a role in their evolution.

Understanding the mechanisms and drivers of macroecological change over geological time scales is a major goal for palaeobiologists who aim to bridge the gap between palaeontology and ecology. Insights may be gained by taking a multi-pronged approach and combining differing diversification dynamics models with environmental data from the geological record. Using large data sets such as these also allows us to feed into the “big data” questions that are asked by biodiversity informaticians. Peterson et al. (2010) identified five big questions for biodiversity informatics; under the approach taken here we are able to contribute to the second of these – a “biota-wide picture of diversification and interactions”. The effects of climate change on the biota are clearly complex and multi-variate. Much data on past climate change and biotic interactions are held within the geological record; and analyses such as these are the first step towards a better understanding of how present day biota might interact and respond to further environmental change. A full global assessment of extinction risk is yet to be carried out for Orthoptera, but the IUCN (2018) has found that more than a quarter of European species are threatened with extinction due to habitat loss. Insects are a vital component of ecosystems, particularly as a food source for vertebrates such as reptiles and birds, and their loss would have cascading ecosystem effects. Combining a macroevolutionary approach to macroecology with present day risk assessments is one way in which we might begin to address another of the aforementioned “big questions” - synthetic conservation planning – where we can integrate knowledge of the past with the present to understand the broader context in which ecosystems and their constituent species evolved, and to help guide conservation efforts.

### Statement of authorship

KED designed the study, carried out the analyses and wrote the manuscript. JH performed the Transfer Entropy analyses. AB & HS contributed data. HS & PM suggested additional analyses. All authors contributed to manuscript revisions.

## Acknowledgements

This work was supported by the Leverhulme Trust, project grant no. RPG-2016–201 awarded to KED, JH and PM. The authors would also like to thank David Chesmore and Edward Baker (University of York) for their respective roles in obtaining the funding that led to this research. Finally, the authors would like to thank Graeme Lloyd (University of Leeds) for comments that helped to improve this manuscript.

